# Endothelial cells elicit a pro-inflammatory response to SARS-CoV-2 without productive viral infection

**DOI:** 10.1101/2021.02.14.431177

**Authors:** Lilian Schimmel, Keng Yih Chew, Claudia Stocks, Teodor Yordanov, Patricia Essebier, Arutha Kulasinghe, James Monkman, Anna Flavia Ribeiro dos Santos Miggiolaro, Caroline Cooper, Lucia de Noronha, Anne K. Lagendijk, Kate Schroder, Larisa I. Labzin, Emma J. Gordon, Kirsty R. Short

**Affiliations:** Division of Cell and Developmental Biology, Institute for Molecular Bioscience, The University of Queensland, Brisbane, QLD, Australia; School of Chemistry and Molecular Biosciences, The University of Queensland, Brisbane, QLD, Australia; IMB Centre for Inflammation and Disease Research, Institute for Molecular Bioscience, The University of Queensland, Brisbane, QLD, Australia; Queensland University of Technology, School of Biomedical Sciences, Faculty of Health, Brisbane, Queensland, Australia; Translational Research Institute, Brisbane, Queensland, Australia; Postgraduate Program of Health Sciences, School of Medicine, Hospital Marcelino Champagnat - Pontifícia Universidade Católica do Paraná (PUCPR), Brazil; Pathology Queensland, Princess Alexandra Hospital, Brisbane, Qld, Australia; School of Medicine & Center of Education, Research and Innovation – Hospital Marcelino Champagnat - Pontifícia Universidade Católica do Paraná (PUCPR). Brazil

**Keywords:** SARS-CoV-2, endothelial cells, COVID-19, inflammation, blood vessels

## Abstract

**Objectives:** Thrombotic and microvascular complications are frequently seen in deceased COVID-19 patients. However, whether this is caused by direct viral infection of the endothelium or inflammation-induced endothelial activation remains highly contentious.

**Methods:** Here, we use patient autopsy samples, primary human endothelial cells and an *in vitro* model of the pulmonary epithelial-endothelial cell barrier to show that primary human endothelial cells express very low levels the SARS-CoV-2 receptor ACE2 and the protease TMPRSS2.

**Results:** Accordingly, endothelial cells can only be infected when SARS-CoV-2 is present at very high concentrations. However, this is not a productive infection (i.e. no infectious virus is produced) and viral entry induces an inflammatory response. We also show that SARS-CoV-2 does not infect endothelial cells in 3D vessels under flow conditions. We further demonstrate that in a co-culture model endothelial cells are not infected with SARS-CoV-2. They do however sense and respond to infection in the adjacent epithelial cells, increasing ICAM-1 expression and releasing pro-inflammatory cytokines.

**Conclusions:** Taken together, these data suggest that *in vivo*, endothelial cells are unlikely to be infected with SARS-CoV-2 and that infection is only likely to occur if the adjacent pulmonary epithelium is denuded (basolateral infection) or a high viral load is present in the blood (apical infection). In such a scenario, whilst SARS-CoV-2 infection of the endothelium can occur, it does not contribute to viral amplification. However, endothelial cells are still likely to play a key role in SARS-CoV-2 pathogenesis by sensing adjacent infection and mounting a pro-inflammatory response to SARS-CoV-2.

## INTRODUCTION

SARS-CoV-2 causes diverse clinical syndromes, ranging from asymptomatic infection to fatal disease. Of patients hospitalised with COVID-19 (the clinical manifestation of SARS-CoV-2 infection), approximately 30% of individuals go on to develop severe disease associated with progressive lung damage ^1^. This is associated with a breakdown of the vascular barrier, oedema, endotheliitis, thrombosis and inflammatory cell infiltration ^1, 2^. Many studies have identified thrombotic and microvascular complications in deceased patients, suggesting that vascular pathology is a major driver of severe disease ^3, 4^. Other major pathological events include arterial and venous thromboembolism, kidney disease and neurological disorders ^5, 6^, suggesting that SARS-CoV-2 activates the vasculature throughout the body, potentially resulting in multi-organ failure. However, whether this is caused by direct viral infection of the endothelium or inflammation-induced endothelial activation remains highly contentious.

SARS-CoV-2 cellular uptake is mediated by binding of the spike glycoprotein to ACE2 (angiotensin-converting enzyme 2) and NRP1 (Neuropilin-1) at the cell surface ^7–10^. Host surface proteases such as TMPRSS2 cleave full-length spike protein (S0) at its S2’ site. Cleavage at the S2’ site can facilitate the fusion of viral and cellular membranes to deliver the viral RNA into the cytosol ^1, 11^. Alternative routes of virus uptake have also been reported in cells with low expression of TMPRSS2: this involves endo-lysosomal pathway and cathepsins ^12^. Numerous studies suggest that endothelial cells express ACE2 ^3, 13–17^, Neuropilin receptors ^18–22^ and TMPRSS2 ^23^, suggesting that viral infection of endothelial cells is theoretically possible. However, others have suggested that ACE2 is highly expressed in microvascular pericytes ^14, 24, 25^ and that pericyte injury in response to viral infection may lead to endothelial dysfunction.

Evidence regarding direct infection of the endothelium with SARS-CoV-2 remains equivocal ^26, 27^. Autopsy studies on COVID-19 patients have suggested the presence of viral particles in the vascular beds of different organs ^3, 28, 29^ and others have shown SARS-CoV-2 infection of cultured endothelial cells ^30, 31^ and human blood vessel organoids ^32^. Autopsy studies of deceased COVID-19 patients have also shown SARS-CoV-2 RNA to be enriched in pulmonary endothelial cells ^33^. In contrast, several studies have failed to find evidence of endothelial infection in deceased COVID-19 patients ^27^ and called into question the validity of identifying virally-infected cells based on the presence of the SARS-CoV-2 spike protein ^25, 34, 35^. Animal models of SARS-CoV-2 infection have also failed to identify any obvious signs of endothelial infection ^36, 37^, despite clear endothelial dysfunction and thrombosis ^38^.

When and how endothelial cells may be exposed to SARS-CoV-2 *in vivo* also remains unclear. As a respiratory virus, SARS-CoV-2 encounters pulmonary epithelial cells prior to any interaction with the endothelium. SARS-CoV-2 infects pulmonary epithelial cells apically and new viral particles are also released apically into the lumen of the lung ^39^. It is possible that the infected epithelial layer loses barrier function, allowing endothelial infection from the basolateral side of epithelia that is adjacent to alveolar endothelium. Alternatively, SARS-CoV-2 may infect endothelial cells apically via the blood, although viremia appears to be rare in COVID-19 patients ^40^. The question therefore remains whether endothelial dysfunction in COVID-19 is the result of direct viral infection of the endothelium.

Here, using patient autopsy samples and *in vitro* models, we show that whilst SARS-CoV-2 can enter endothelial cells if high viral titres are present, this infection is not productive. Rather, in response to direct or indirect viral exposure, endothelial cells mount a pro-inflammatory response characterised by increased expression of ICAM-1 and secretion of CXCL-10 and IL-6. Taken together, our results provide clarification on the intensely debated topic of endothelial infection by SARS-CoV-2, to reveal that the endotheliopathy and thrombocytopathy in patients with severe COVID-19 is likely to be due to the inflammatory response, rather than direct viral infection.

## MATERIALS AND METHODS

### In vivo samples

Lung tissue from 10 SARS-CoV-2 infected patients was sectioned and Hematoxylin and Eosin (H&E) staining performed. RNAscope probes (ACDbio, US) targeting SARS-CoV-2 spike mRNA (#848561-C3), were used as per manufacturer instructions for automation on Leica Bond RX. DNA was visualised with Syto13, and SARS-CoV-2 spike probe with opal 690 (1:1500). Fluorescent images were acquired with Nanostring Mars prototype DSP at 20x. Autopsy and biopsy materials were obtained from the Pontificia Universidade Catolica do Parana PUCPR the National Commission for Research Ethics (CONEP) under ethics approval numbers 2020001792/30188020.7.1001.0020 and 2020001934/30822820.8.000.0020. The study was also approved under University of Queensland HREC ratification.

### Cell culture

Human umbilical vein endothelial cells (HUVECs), and human microvascular endothelial cells from lungs (HMVEC-L) purchased from Lonza (Cat# CC-2935, and CC-2527 respectively) were cultured until passage 8 in EGM-Plus or EGM-2MV medium, supplemented with singlequots (Lonza Cat# CC-5035, CC-3102). Calu-3 cells purchased from ATCC (Cat# HTB-55) were maintained in MEM (Invitrogen), containing 10% (v/v) heat-inactivated foetal bovine serum (Cytiva), 100 U/ml penicillin and streptomycin (Life Technologies Australia), and grown in EGM-2MV for endothelial co-culture experiments. Baby hamster kidney cells (BHK-21) were a kind gift from Prof. Dr. R.G. Parton. Cell lysates of human nasal epithelial cells (HNEp) were a kind gift from Prof. Peter Sly.

Monocultures of HUVEC, HMVEC or Calu-3 cells were performed on 24 well cell culture inserts (Corning 6.5 mm Transwell, 0.4 μm polycarbonate membrane Cat#3413) coated with 5 μg/ml fibronectin (FN) (Sigma). Approximately 25,000 HUVEC, 25,000 HMVEC and 50,000 Calu-3 cells were seeded per insert on either the apical or basolateral side as indicated and grown in EGM-Plus (Lonza), EGM-2MV (Lonza) or MEM (Invitrogen) respectively. Endothelial cells were treated with 100ng/ml TNFα (Life Technologies Australia) or 10ng/ml IFNß (Lonza) for as long as indicated.

Calu-3 and HMVEC were co-cultured on 24 well cell culture inserts (Corning 6.5 mm Transwell, 0.4 μm polycarbonate membrane Cat#3413) coated with 5 μg/ml FN (Sigma) in EGM-2MV (Lonza). Approximately 25,000 HMVECs were seeded on an inverted transwell membrane for 1 hr to attach. Transwell membranes were turned over to normal position and approximately 50,000 Calu-3 cells were seeded in the top compartment of the membrane. Cells were grown for 48 hr prior to infections.

### Viral infection

SARS-CoV-2 isolate hCoV-19/Australia/QLD02/2020 was provided by Queensland Health Forensic & Scientific Services, Queensland Department of Health. Virus was grown on Vero cells and titred ^41^. Sanger sequencing was used to confirm that no mutations occurred in the spike gene relative to the original clinical isolate. Cells were infected with 6 x 10^4^ plaque forming units (PFU) for 1 hour at 37 degrees 5% Co_2_. The viral inoculum was then removed, and the medium was replaced with DMEM (Invitrogen) or MEM (Invitrogen) containing 2% FBS. Alternatively, 2 x 10^6^ PFUs were added to the cell culture and cells were incubated at 37 degrees 5% CO2. All studies with SARS-CoV-2 were performed under physical containment 3 (PC3) conditions and were approved by the University of Queensland Biosafety Committee (IBC/374B/SCMB/2020).

### Antibodies

The following antibodies were used: mouse anti-ACE2 (Santa Cruz, sc-390851, IF1:200) goat anti-ACE2 (R&D systems, AF933, WB 1:1000), mouse anti-dsRNA (Millipore, MABE1134, IF1:100), rabbit anti-SARS-CoV nucleocapsid protein/NP (Sino Biological, 40143-R040, WB1:1000 IF1:200), rabbit anti-TMPRSS2 (Abcam, ab92323, WB1:1000), rabbit anti-cleaved caspase 3 (Cell Signaling, 9661 IF1:30 after pre-labeling), rabbit anti-GAPDH (Cell Signaling, 2118, WB1:2000), mouse anti-ICAM-1 (R&D systems, BBA3, IF1:200), mouse anti-ICAM-1 (Santa Cruz Biotechnology, sc-8439, WB1:1000), Phalloidin-Alexa555 (Cytoskeleton, PHDH1-A, IF1:500), Phalloidin-Alexa670 (Cytoskeleton, PHDN1-A, IF1:200). Pre-labeling of rabbit anti-cleaved caspase 3 with Alexa Fluor 555 was performed using Zenon Rabbit IgG labelling kit (Z-25305) according to manufacturer protocol.

### Immunofluorescence Staining

Immunofluorescence staining was in general performed on cells cultured on 12 mm glass coverslips or on 24 well cell culture inserts (Corning 6.5 mm Transwell, 0.4 μm polycarbonate membrane Cat#3413) coated with 5 μg/ml FN (Sigma), washed with PBS^+/+^(PBS supplemented with 1 mM CaCl_2_ and 0.5 mM MgCl_2_), fixed for 10 min in 4% PFA (Sigma), blocked and permeabilized for 30 min with 3% BSA, 0.3% Triton X-100 (Sigma). Primary antibodies were incubated in 1.5% BSA for 1 hr at RT, secondary antibodies linked to Alexa fluorophores (all Invitrogen) were incubated in 1.5% BSA for 1 hr at RT and incubated with 4% PFA (Sigma) for another 24 hr in case of SARS-CoV-2 infected cells, after each step samples were washed 3x with PBS^+/+^ and mounted in ProlongGold + DAPI solution (Cell Signaling Technologies, CST 8961S). Z-stack image acquisition was performed on a confocal laser scanning microscope (Zeiss LSM880) using a 40x NA 1.3 or 63x NA 1.4 oil immersion objective.

### 3D Microfluidic cell culture

Microfluidic devices were generated as previously described ^42^ using 2.5mg/ml Collagen (R&D Systems) for the ECM. Cells were seeded 48 hr prior to virus infection and were kept on a rocker at 37C° and 5%CO_2_. The tubes were fixed sequentially in 1% formaldehyde, containing 0.05% Triton for 90 sec, followed by 4% formaldehyde for 15 min at 37C°. Tubes were permeabilised with 0.5% Triton for 10 min at 37C° and blocked for 4 hr in 10% goat serum in PBS at 4C° prior to staining. Immunofluorescence staining was performed using NP, dsRNA and Phalloidin-Alexa670 antibodies. Z-stack image acquisition was performed on a confocal laser scanning microscope (Zeiss Axiovert 200 inverted microscope with LSM 710 Meta Confocal Scanner) using 40X NA 1.1 water immersion objective.

### Western blotting

For total cell lysates, cells were washed once with PBS and lysed with RIPA buffer (50 mM Tris, 150 mM NaCl, 1 mM EDTA, 1% Triton X-100, 0.1% SDS, 1% sodium deoxycholate, protease inhibitor, pH 8.0). Pierce BCA protein assay kit (Thermo Scientific) was used to equalize protein amounts and SDS-sample buffer containing 100 mM DTT (Astral Scientific) was added and samples were boiled at 95°C for 10 minutes to denature proteins. Proteins were separated on 4-15% mini protean TGX precast gels (Biorad) in running buffer (200 mM Glycine, 25 mM Tris, 0.1% SDS (pH8.6)), transferred to nitrocellulose membrane (BioRad Cat#1620112) in blot buffer (48 nM Tris, 39 nM Glycine, 0.04% SDS, 20% MeOH) and subsequently blocked with 5% (w/v) BSA in Tris-buffered saline with Tween 20 (TBST) for 30 minutes. The immunoblots were analysed using primary antibodies incubated overnight at 4°C and secondary antibodies linked to horseradish peroxidase (HRP) (Invitrogen), and after each step immunoblots were washed 4x with TBST. HRP signals were visualized by enhanced chemiluminescence (ECL) (BioRad) and imaged with a Chemidoc (BioRad).

### RNA extraction and cDNA synthesis

RNA was extracted using the Qiagen RNeasy Minikit (Qiagen) or NucleoZOL (BIOKÉ, The Netherlands) according to manufacturer’s guidelines. cDNA was synthesized using random hexamers and the high-capacity cDNA reverse transcription kit (Life Technologies) according to the manufacturer’s guidelines.

### Quantitative PCR (qPCR)

qPCR was performed using SYBR Green reagent (Applied Biosystems). All primer sequences are listed in Table 1. qPCR conditions were applied according to the manufacturer’s instructions using QuantStudio 6 Flex Real-Time PCR System (ThermoFisher Scientific, Waltham, Massachusetts, MA). HPRT or Glyceraldehyde 3-phosphate dehydrogenase (GAPDH) was used as a housekeeping gene and relative gene expression was determined using the comparative cycle threshold (ΔΔCt) method (Livak, 2001).

**Table 1:**
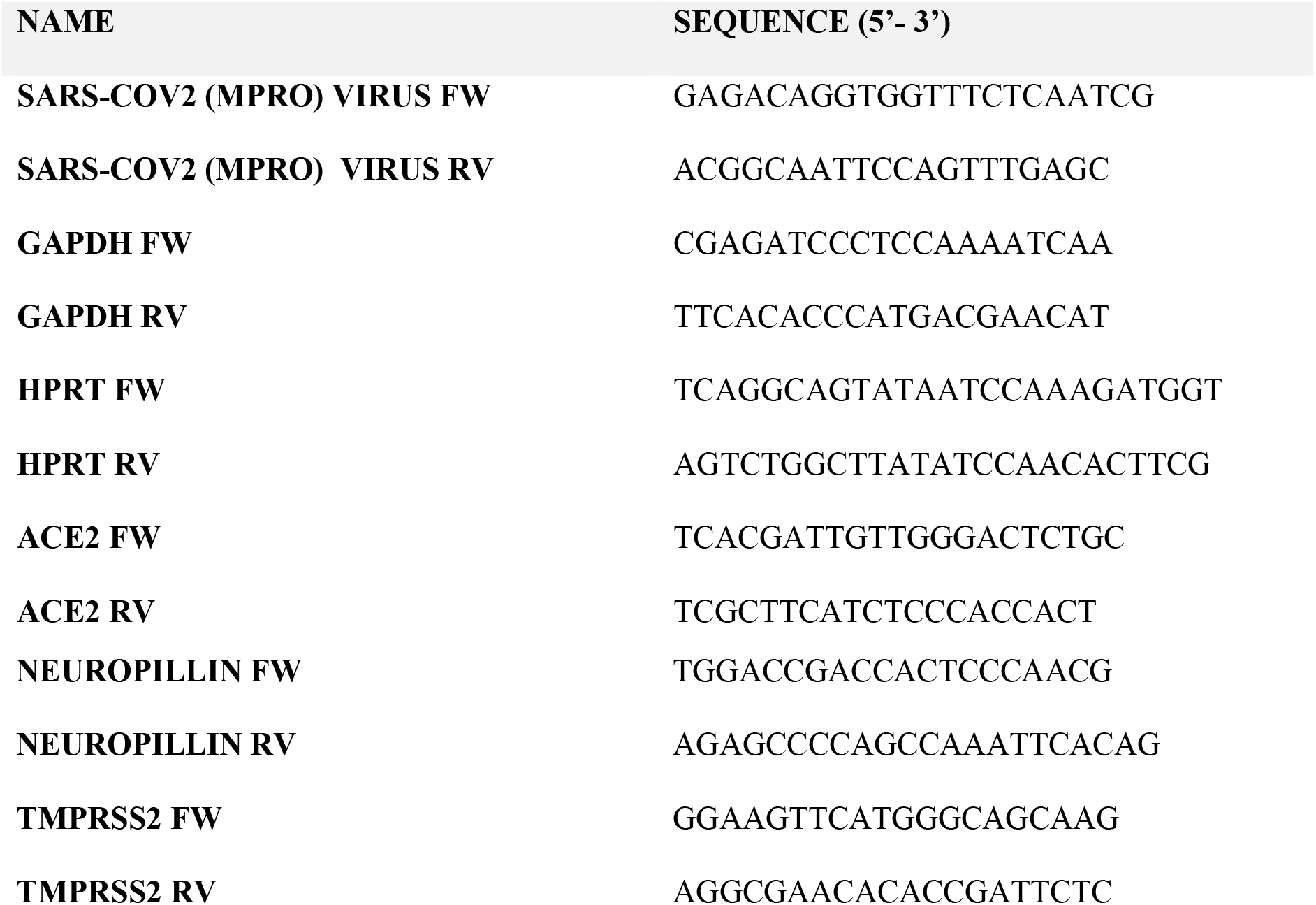
Primers used in the present study

### Viral titres

Viral titres in cell culture supernatant determined by a plaque assay on Vero cells, according to previously described protocols ^41^.

### Cytokine titres

Cytokine titres in were determined using an Alphalisa kit (Perkin Elmer) according to the manufacturer’s instructions. The sum of cytokines detected in the upper and lower compartments of the transwell is shown for each of the monocultured samples, while cytokine levels were kept separated for the upper and lower compartment for the co-cultured samples.

### Image analysis

Analysis of immunofluorescent images was performed using ImageJ version 2.0.0-rc-69/1.52n. A mask of the total cell area was created using default intensity threshold on Phalloidin staining. Within the mask, Raw Integrated Density of either ACE2, NP, ICAM-1 was measured, and value was corrected for total mask area resulting in RawIntDen/Area. For dsRNA, MaxEntropy threshold was performed on dsRNA staining to remove background speckles before measuring Raw Integrated Density within the mask. Analysis of Western blot images was performed using Gel Analyzer Tool of ImageJ version 2.0.0-rc-69/1.52n. The background signal was subtracted and values were normalized to corresponding GAPDH loading control. Imaris (Bitplane) version 8 was used to create the XZ projection in Figure 4B”. Extrusion of cells was quantified in Supplemental Figure 1C using orthogonal views in ImageJ version 2.0.0-rc-69/1.52n.

### Statistical analysis

All statistical analysis was performed using Graphpad Prism version 9.0. Data are presented as mean±s.e.m. with individual data points indicated and colour coded per independent experimental replicate. Statistical significance was determined using Kruskal Wallis test for multiple comparisons or Mann-Whitney test between mock and SARS-CoV-2 treated conditions. *P<0.05, **P<0.01, ***P<0.001 and ****P<0.0001.

## RESULTS

### Endothelial cell infection is not readily detected in deceased COVID-19 patients

To determine the prevalence of *in vivo* SARS-CoV-2 endothelial cell infection, autopsy lung sections were obtained from 10 deceased COVID-19 patients and probed for SARS-CoV-2 spike mRNA using RNAscope. Two of the 10 patients were positive for SARS-CoV-2 RNA in the lungs. In these individuals, spike mRNA could not be detected in pulmonary endothelial cells (endothelial cells defined by H & E staining (Figure 1). These data suggest that the endothelium is not generally the primary site of viral replication *in vivo* in COVID-19 patients (patient data available in Supplemental Table 1).

**Figure 1.**
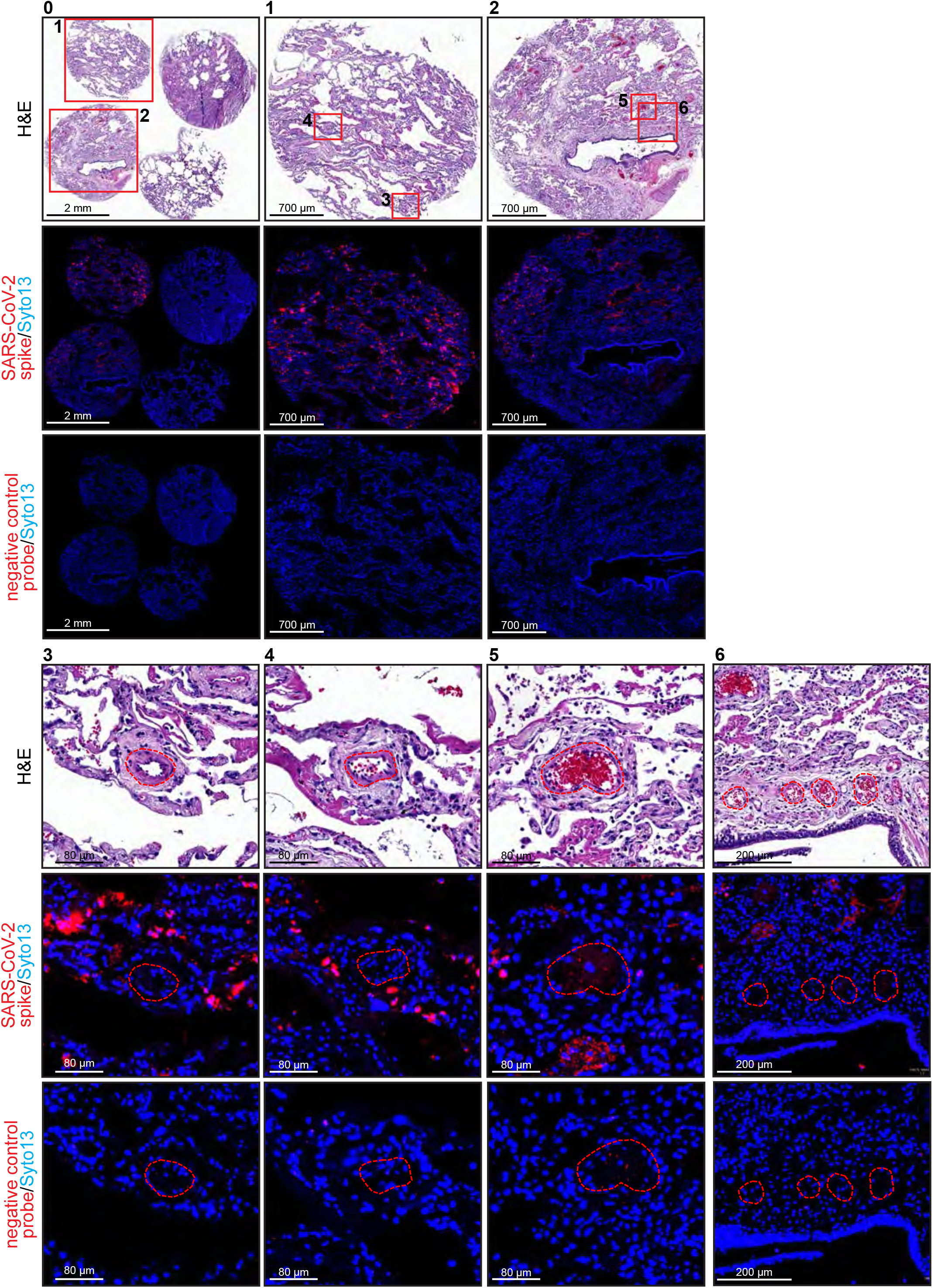
In vivo sections of lung show endothelial cells are not infected. Lung tissue from 4 deceased COVID-19 patients was sectioned and stained for Hematoxylin and Eosin (H&E)(0). Consecutive sections were stained using RNAscope probes targeting either SARS-CoV-2 spike mRNA or negative control probe and Syto13 for DNA. Patients 1 and 2 are positive for SARS-CoV-2 spike mRNA (enlargement 1 and 2). Further enlargements of numbered insets (3-6) show vessels outlined with red dashed line are negative for SARS-CoV-2 spike mRNA in the endothelial layer of the vessels. Negative control probe shows non-specific background staining.

### Primary endothelial cells express low levels of ACE2 and TMPRSS2

To further investigate the contribution of endothelial cells to the pathogenesis of SARS-CoV-2, we established *in vitro* cultures of primary human umbilical vein endothelial cells (HUVECs) and human microvascular endothelial cells from the lung (HMVEC-L). Multiple studies have reported that endothelial cells across different vascular beds express ACE2 and TMPRSS2, the host cell receptor and protease that are required for efficient cell infection by SARS-CoV-2 ^10, 11^. We found that both HUVECs and HMVEC-Ls expressed comparable ACE2 and TMPRSS-2 mRNA to that observed in immortalised epithelial cells known to be susceptible to SARS-CoV-2 infection (Calu-3) (Figure 2A). Neuropilin-1, which enhances viral entry through its interaction with the spike multibasic cleavage site ^7, 8^, was readily detected in endothelial cells by qPCR (Figure 2A) in line with established data ^19^. The expression of ACE2 protein detected by immunofluorescence (Figure 2B) was significantly reduced compared to epithelial cells (Calu-3) (Figure 2C). We also detected ACE2 and TMPRSS2 protein in HUVECs and HMVEC-Ls by western blot (Figure 2D-E). The specificity of our ACE2 antibody was confirmed by western blot of BHK-21 cells (which are known to be negative for ACE2) ^43^.These data suggest that while vascular endothelial cells can theoretically be infected by SARS-CoV-2, infection is likely to be inefficient compared to epithelial cells, because of weak receptor and protease expression in endothelial cells.

**Figure 2.**
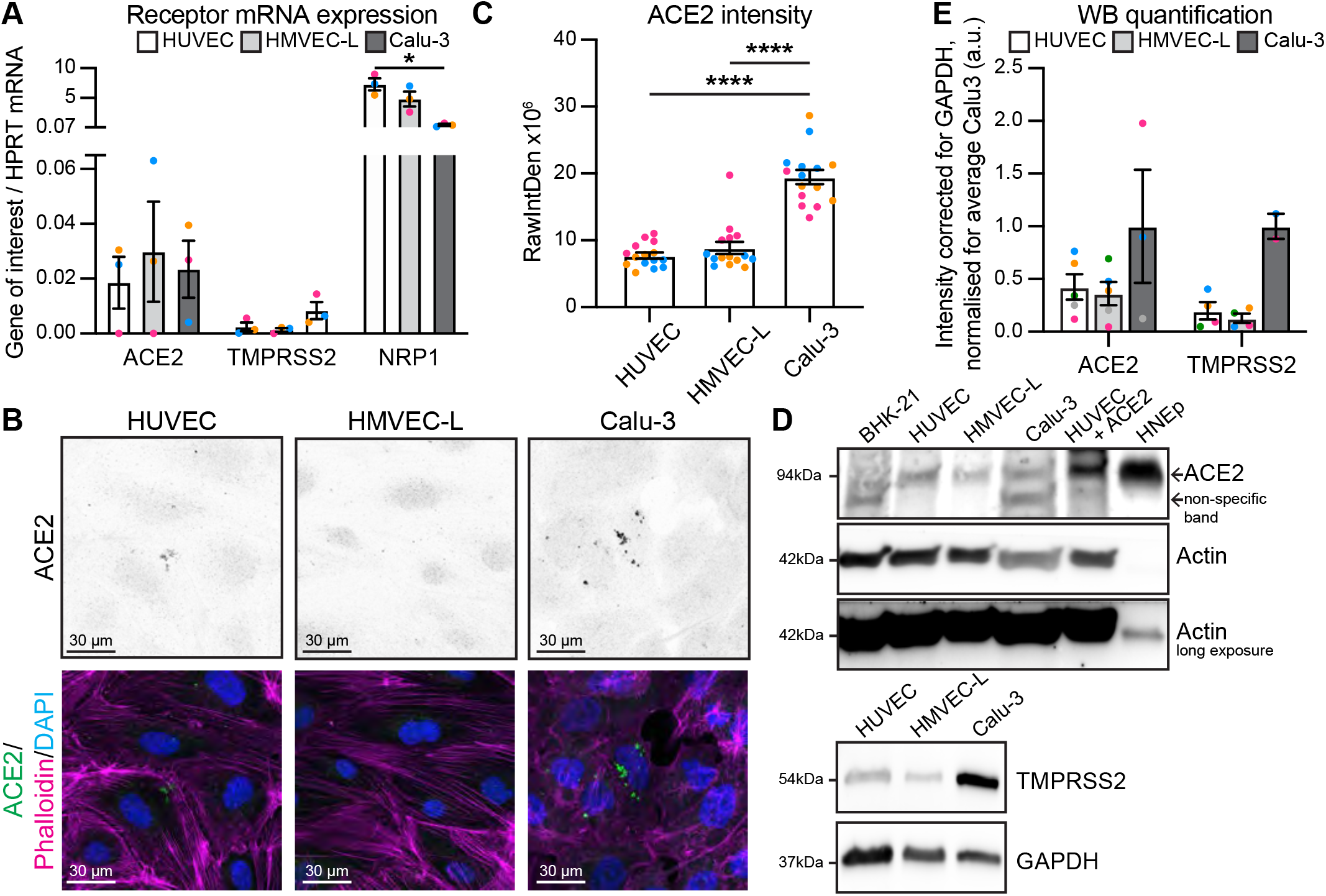
Endothelial cells express very low levels of ACE2 and TMPRSS2 receptors. **A)** qPCR shows presence of mRNA for ACE2, TMPRSS2 and NRP1 in HUVEC, HMVEC-L and Calu-3 cells. n=3 independent experiments. **B)** Representative immunofluorescent images of HUVEC, HMVEC-L and Calu-3 cells stained for ACE2 (shown as single channel in top panel) (green), Phalloidin (magenta) and DAPI (blue). Scalebar 30 μm. **C)** Quantification of ACE2 staining intensity. n=15 images; 5 images per independent experiments. **D)** Western blot analysis showing ACE2 protein levels are absent in negative control BHK-21 cells, lower in HUVEC and HMVEC-L compared to Calu-3 cells and HUVEC + ACE2 overexpression and Human Nasal Epithelium cells (HNEp) as positive controls in top panel. TMPRSS2 protein levels are lower in HUVEC and HMVEC-L compared to Calu-3 cells in bottom panel **E)** Quantification of protein levels for ACE2 and TMPRSS2 in HUVEC, HMVEC-L and Calu-3. n=5 (HUVEC and HMVEC-L), n=3 (Calu-3) independent experiments. **D)** Data are presented as mean±s.e.m. with individual data points indicated and colour coded per independent experimental replicate. Statistical significance was determined using a Kruskal Wallis test between Calu-3 and all others (A, C, E). *P<0.05, ****P<0.0001.

### Endothelial cells are not productively infected by SARS-CoV-2

SARS-CoV-2 may theoretically enter endothelial cells via the apical surface (should the virus enter the blood stream) or the basolateral surface (should the virus be present at the basolateral surface of the adjacent epithelial cells). We therefore assessed whether endothelial cells were susceptible to SARS-CoV-2 infection when exposed either apically (Supplemental Figure 1A) or basolaterally (Supplemental Figure 1B) to 6 x 10^4^ PFUs. Infectious virus was not detected in the supernatant of either HUVECs or HMVEC-Ls at 24, 48 nor 72 hours post-infection (Supplemental Figure 1C). In contrast, Calu-3 cells demonstrated robust viral replication and release (Supplemental Figure 1C). Similarly, no increase in SARS-CoV-2 nucleocapsid protein within infected endothelial cells relative to mock was detected by immunofluorescence (Supplemental Figure 1D-G) or western blot (Supplemental Figure 1H-I), and no detectable alterations in endothelial cell morphology were evident after viral exposure, as assessed by phalloidin immunostaining (Supplemental Figure 1D, F). In contrast, Calu-3 cells displayed a significant increase in nucleocapsid protein expression after viral exposure. Together, these data suggest that endothelial cells *in vitro* are not efficiently infected with SARS-CoV-2.

We next sought to establish if the low levels of ACE2 and TMPRSS2 on endothelial cells could be overcome by a higher viral titre, to mediate infection. Accordingly, endothelial cells were infected either apically or basolaterally with 2 x 10^6^ PFUs of SARS-CoV-2 and viral replication was again assessed over time. Infectious virus did not increase over time in endothelial cells, rather viral titres decreased over time, consistent with a degradation of input virus (Figure 3A). In contrast, productive viral replication was observed in Calu-3 cells over time (Figure 3A, Supplemental Figure 2A), while, as expected, non-susceptible, ACE2-negative BHK-21 cells showed no productive viral replication (Supplemental Figure 2A). To determine if endothelial cells were susceptible to viral entry, if not replication, we assessed viral nucleocapsid protein by immunofluorescence in HMVEC-L (Figure 3B-C) positive control Calu-3 and negative control BHK-21 (Supplemental Figure 2D-E) and western blot (Figure 3E-F, Supplemental Figure 2B-C). Viral nucleocapsid protein expression was significantly increased after both apical and basolateral SARS-CoV-2 infection, revealing that SARS-CoV2 can indeed enter endothelial cells *in vitro*. We observed that the majority of infected, nucleocapsid protein-positive endothelial cells appeared to be extruding from the cell monolayer in an apical manner (Figure 3B’, 3B”, 3C). These results were not due to dead cells present in the viral inoculum as BHK-21 cells did not show any nucleocapsid-positive extruded cells (Supplemental Figure 2D). Moreover, extruding endothelial cells were positive for cleaved caspase 3, indicating that SARS-CoV-2 entry into endothelial cells induces apoptosis (Figure 3G-H). These results demonstrate that while SARS-CoV-2 can enter endothelial cells this infection is not productive and leads to apoptosis and extrusion of these apoptotic cells from the endothelial monolayer.

**Figure 3.**
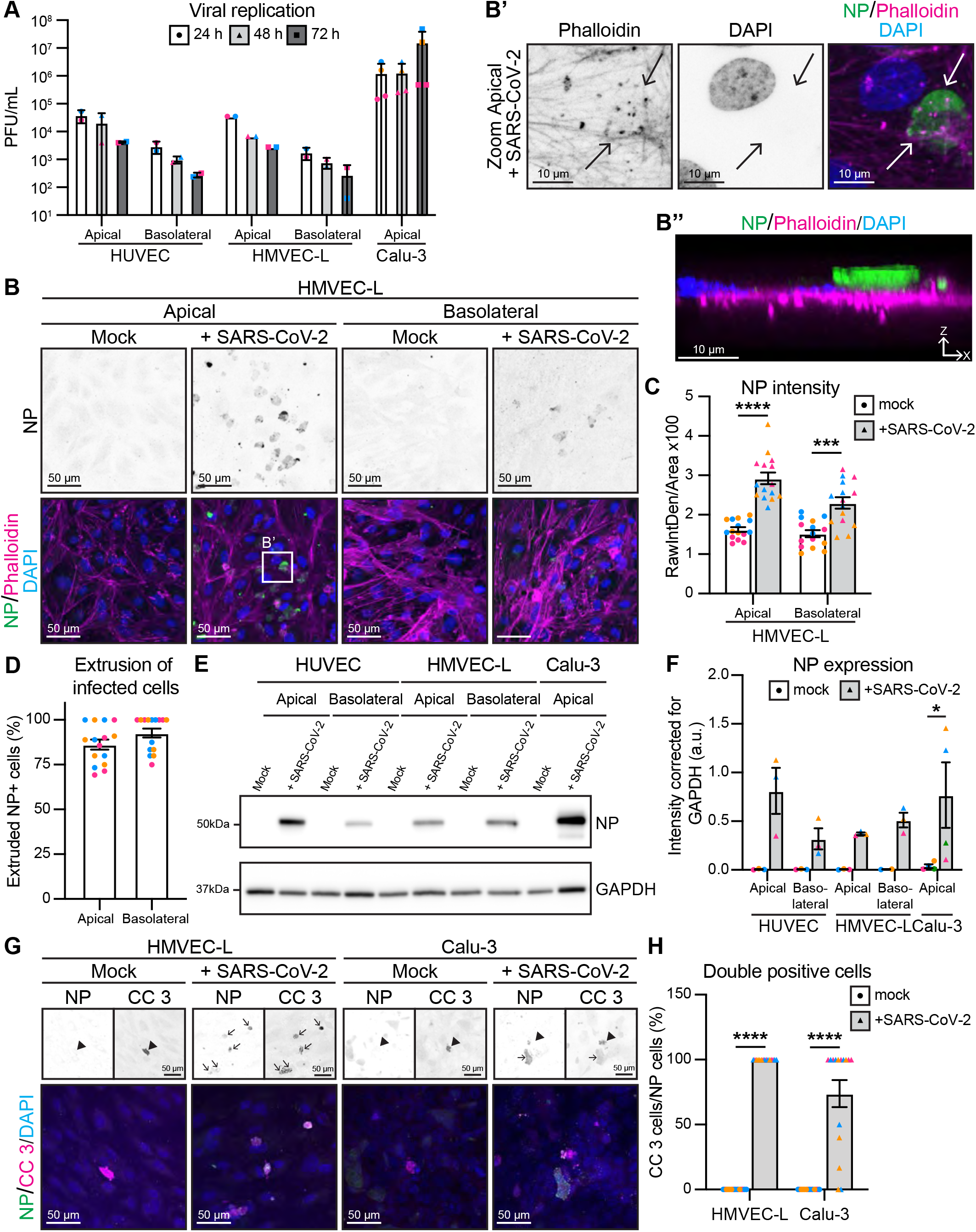
Endothelial cells can be infected with 2 x 10^6^ PFU of SARS-CoV-2, but infection is abortive. **A)** Viral replication shown as number of PFU per ml of supernatant from SARS-CoV-2 infected HUVEC, HMVEC-L and Calu-3 cells at 24h, 48h and 72h after infection. n=2 (HUVEC and HMVEC-L), n=3 (Calu-3) independent experiments. **B)** Representative immunofluorescent images of HMVEC-L stained for NP (shown as single channel in top panel) (green), phalloidin (magenta) and DAPI (blue) with mock or SARS-CoV-2 infection from either apical or basolateral side of the cells at 72h after infection. Scalebar 50 μm. **B’)** Enlargement of white box in B shows Phalloidin and DAPI single channels and merge. Scalebar 10 μm. **B’’)** XZ projection of merged image in B’, with apical side on top of image and basolateral side on the bottom. Scalebar 10 μm. **C)** Quantification of NP staining intensity in HMVEC-L. n=15 images from 3 independent experiments. **D)** Quantification of % NP positive cells that are extruded upon SARS-CoV-2 infection of HMVEC-Ls. n=15 images from 3 independent experiments. **E)** Western blot analysis showing NP protein levels in HUVEC, HMVEC-L and Calu-3 cells after 72h of infection. **F)** Quantification of NP protein levels in HUVEC, HMVEC-L and Calu-3. n=3 independent experiments. **G)** Representative immunofluorescent images of HMVEC-L and Calu-3 stained for NP (shown as single channel in top left panel) (green), cleaved caspase 3 (CC 3) (shown as single channel in top right panel)(magenta) and DAPI (blue) with mock or SARS-CoV-2 infection from apical side of the cells at 72h after infection. Scalebar 50 μm. **H)** Quantification of % NP positive cells that are positive for apoptosis marker cleaved caspase 3. n=15 images from 3 independent experiments. Data are presented as mean±s.e.m. with individual data points indicated and colour coded per independent experimental replicate. Statistical significance was determined using Kruskal Wallis test between 24h and other time points (A) or Mann-Whitney test between mock and + SARS-CoV-2 (C, F, H). *P<0.05, ***P<0.001, ****P<0.0001.

### Endothelial cells are not productively infected by SARS-CoV-2 in 3D vessels

To determine whether endothelial cells can be infected in 3D vessels under flow, we cultured both HMVEC-L and Vero cells in microfabricated tubes ^42^ and added SARS-CoV2 to the luminal surface (Figure 4A). After 24h, Vero cells were readily infected and viral replication was observed, in contrast to HMVEC-L which shows viral input levels (Figure 4B). In agreement, NP was not detected in HMVEC-L tubes, whereas Vero cells displayed clear NP staining at the luminal surface (Figure 4C-F). This data reveals that fluid flow over endothelial cells does not alter the ability of SARS-CoV-2 to infect cells.

**Figure 4.**
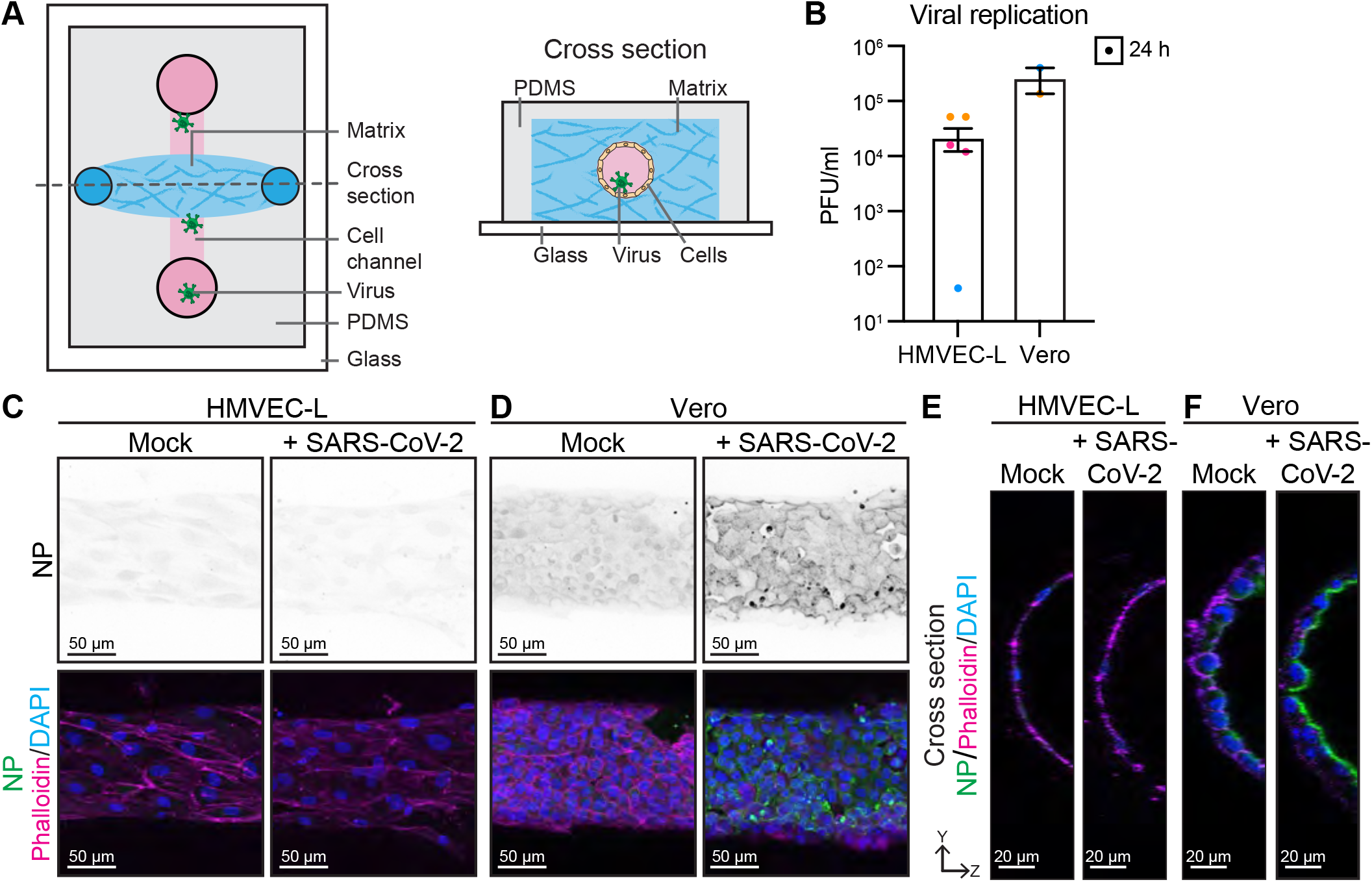
Endothelial cells are not infected with 2 x 10^6^ PFU of SARS-CoV-2 in 3D. **A)** Schematic of microfabricated device used to culture 3D vessel tubes under oscillatory flow. Cross section shows luminized tube covered with endothelial cells surrounded by extracellular matrix and virus can be flowed through the lumen of those tubes. **B)** Viral replication shown as number of PFU per ml of supernatant from SARS-CoV-2 infected HMVEC-L and Vero cells at 24h after infection. n=3 (HMVEC-L), n=2 (Ver0) independent experiments. Representative immunofluorescent images of **C)** HMVEC-L and **D)** Vero stained for NP (shown as single channel in top panel) (green), phalloidin (magenta) and DAPI (blue) with mock or SARS-CoV-2 infection in the lumen of the tube at 24h after infection. Scalebar 50 μm. YZ projection of **E)** HMVEC-L (merged image in C) and **F)** Vero (merged image in D), with luminal side towards the right of image and basolateral side towards the right of images. Scalebar 20 μm.

### Endothelial cells mount a modest immune response to SARS-CoV-2

As severe COVID-19 disease is associated with an elevated cytokine response ^44^, we sought to assess whether the abortive SARS-CoV-2 infection observed in endothelial cells induces a pro-inflammatory response. We assessed expression of the leukocyte adhesion molecule, ICAM-1, which is well established to be induced under inflammatory conditions in the endothelium ^45^. Immunofluorescence showed that HMVEC-L exposed both basally and apically to SARS-CoV-2 significantly increased ICAM-1 expression, except for in nucleocapsid protein-positive HMVEC-L (likely as they are apoptotic and extruded from the monolayer) (Figure 5A-C). The moderate basal ICAM-1 expression observed in mock infected cells is likely due to the fact that the cells are kept for 72h without a media change (as seen in PBS controls (Figure 5B). However, this response was not as robust as when HMVEC-L were exposed to the established inflammatory cytokine TNFα (Figure 5B-C). Western blot analysis confirmed that HMVEC-L exposed to SARS-CoV-2 displayed a trending increase in ICAM-1 expression (Figure 5D, E). In addition to upregulating inflammatory adhesion molecules, we analysed whether endothelial cells can be the source of pro-inflammatory cytokines during SARS-CoV-2 infection. As IL-6 and CXCL10 are elevated in patients with severe COVID-19 ^46^, we analysed whether endothelial cells released these pro-inflammatory cytokines when infected either apically or basolaterally with SARS-CoV-2 for 72h. HMVEC-Ls released modest levels of CXCL10 (Figure 5F) and IL-6 (Figure 5G). While these modest levels of cytokine release from endothelial cells upon SARS-CoV-2 infection are lower than CXCL10 or IL-6 induced in response to high concentrations of recombinant TNF or IFNß, (Supplemental Figure 2F-G), this shows that endothelial cells express high basal levels of IL-6 and that both cytokines are increased during infection. Taken together, these results show that endothelial cells do not support a productive SARS-CoV-2 infection but still mount a modest pro-inflammatory response to the virus.

**Figure 5.**
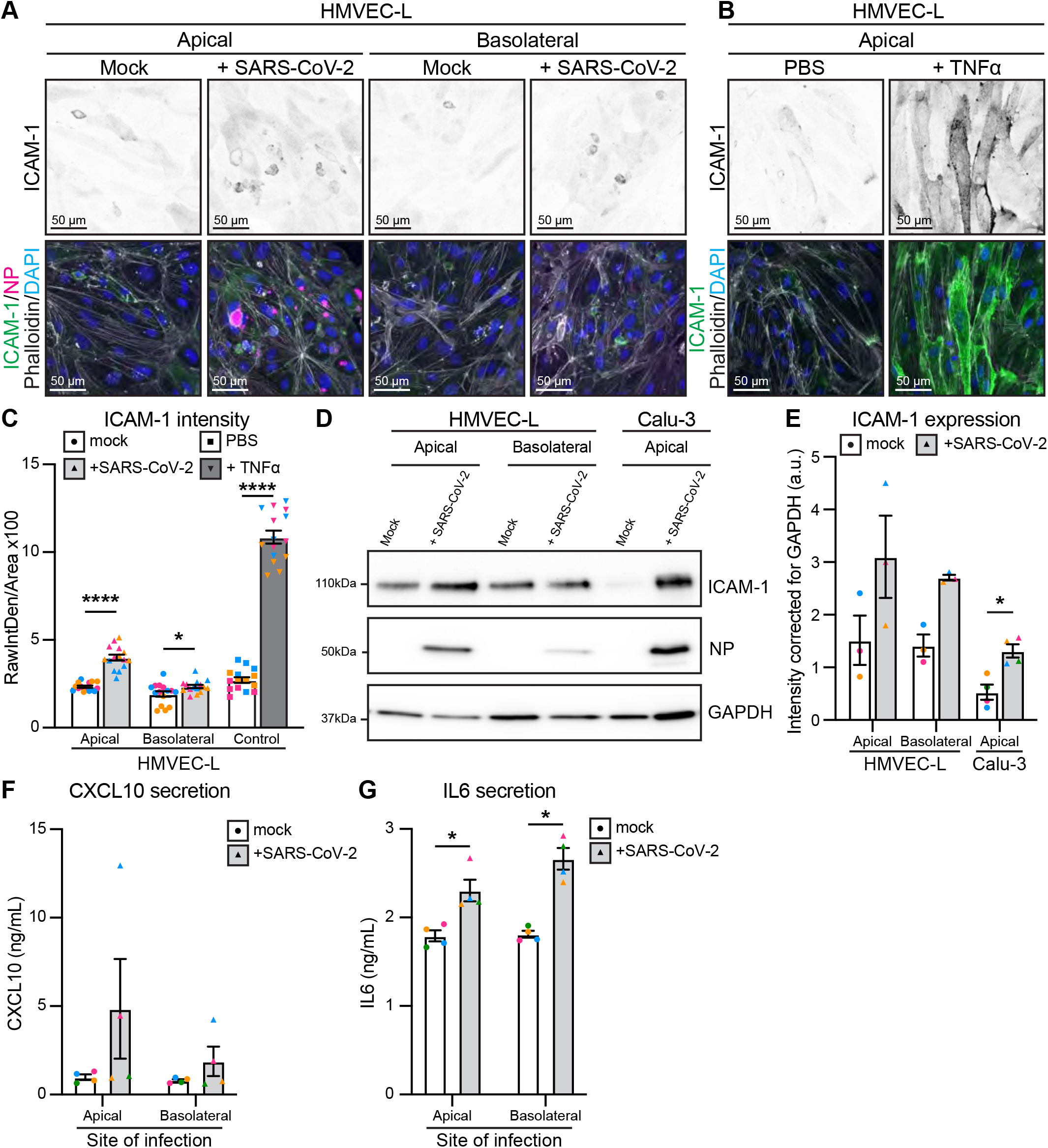
Endothelial cells mount an inflammatory response upon infection. Representative immunofluorescent images of HMVEC-L stained for ICAM-1 (shown as single channel in top panel) (green), NP (magenta), phalloidin (grey) and DAPI (blue) with **A)** mock or SARS-CoV-2 infection from either apical or basolateral side of the cells at 72h after infection or with **B)** control and TNFα treatment for 72h. Scalebar 50 μm. **C)** Quantification of ICAM-1 staining intensity in HMVEC-L. n=15 images from 3 independent experiments. **D)** Western blot analysis showing ICAM-1 and NP protein levels in HMVEC-L and Calu-3 cells after 72h of infection. **E)** Quantification of ICAM-1 protein levels in HMVEC-L and Calu-3. n=3 independent experiments. Measurement of cytokines with AlphaLISA Immunoassay kit for **F)** CXCL10 and **G)** IL6 in the supernatant of HMVEC-L with mock or SARS-CoV-2 infection from either apical or basolateral side of the cells at 72h after infection. n=4 independent experiments. Data are presented as mean±s.e.m. with individual data points indicated and colour coded per independent experimental replicate. Statistical significance was determined using Mann-Whitney test between mock and + SARS-CoV-2 (C, E, F, G). *P<0.05, ****P<0.0001.

### Infection of epithelial-endothelial co-cultures elicits an immune response in endothelial cells

To create a more physiologically relevant *in vitro* model of the pulmonary endothelium, we adapted our previously described co-culture model of the lung epithelial-endothelial barrier ^47^, where epithelial (Calu-3) cells are seeded on top of a transwell membrane while endothelial cells (HMVEC-L) are seeded on the basolateral side (Figure 6C). To mirror a respiratory infection, SARS-CoV-2 was added to the upper compartment and both epithelial and endothelial cells were analysed at 72h post-infection. In contrast to our observations in endothelial cell monocultures, no viral nucleocapsid protein or viral dsRNA was detected in HMVEC-L, whereas SARS-CoV-2 nucleocapsid protein and viral dsRNA (indicating actively replicating virus) was clearly detected in infected Calu-3 cells (Figure 6A,B; 6E,F). These results were confirmed by the presence of higher levels of viral RNA as measured by qPCR in the Calu-3 compared to the HMVEC-L (Fig 6G) and higher titres of infectious virions in the supernatants of the upper (Calu-3) compartment of the transwell (Figure 6H). Despite lack of endothelial infection, we observed that HMVEC-L cells expressed ICAM-1 expression when virus was added to the upper (Calu-3) compartment, suggesting that the HMVEC-L respond to adjacent epithelial infection (Figure 6A, D). These HMVEC-L also responded to the infection in the adjacent Calu-3 cells by increasing CXCL10 secretion (detected in lower compartment of the co-culture) (Figure 6I). Both Calu-3 and HMVEC-L appeared to secrete IL-6 in response to SARS-CoV-2 infection, as IL-6 levels were increased in both compartments (significantly in the upper, trending in the lower) (Figure 6J). In summary, in a co-culture model of the pulmonary epithelium, endothelial cells are not directly infected with the virus, but respond to adjacent epithelial infection by upregulating ICAM-1 expression and releasing pro-inflammatory cytokines. This suggests that endothelial inflammation and dysfunction during *in vivo* SARS-CoV-2 infection is most likely to occur in response to infection of the adjacent epithelial layer.

**Figure 6.**
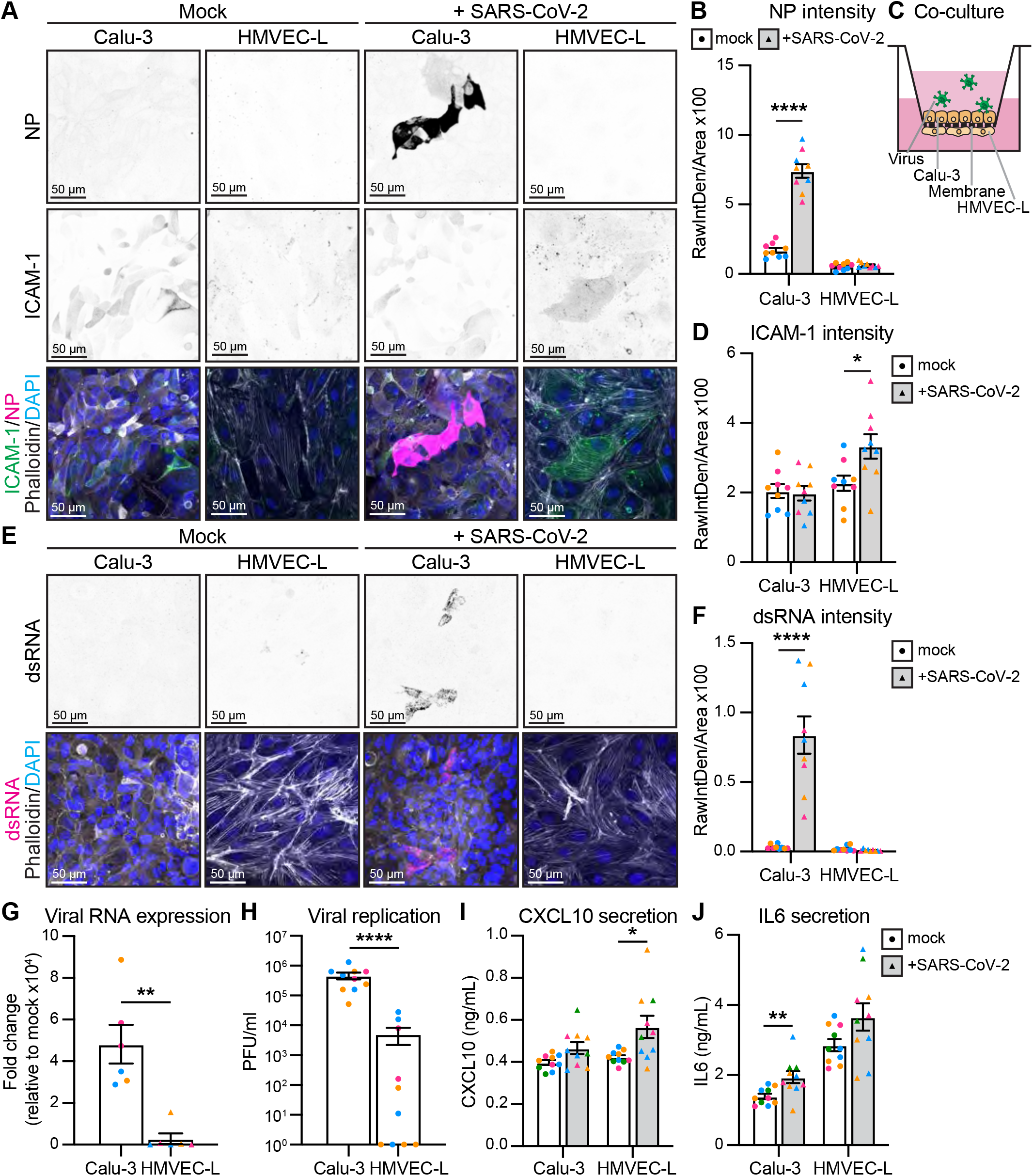
Co-culture shows infection of Calu-3 and inflammation of HMVEC-L. **A)** Representative immunofluorescent images of co-cultured Calu-3 and HMVEC-L stained for NP (shown as single channel in top panel) (magenta), ICAM-1 (shown as single channel in middle panel) (green), phalloidin (grey) and DAPI (blue) with mock or SARS-CoV-2 infection at 72h after infection (2 x 10^6^ PFU). Scalebar 50 μm. **B)** Quantification of NP staining intensity in Calu-3 and HMVEC-L. n=9 images from 3 independent experiments. **C)** Schematic of Calu-3 and HMVEC-L co-cultures on transwell membranes. **D)** Quantification of ICAM-1 staining intensity in Calu-3 and HMVEC-L. n=9 images from 3 independent experiments. **E)** Representative immunofluorescent images of co-cultured Calu-3 and HMVEC-L stained for dsRNA (shown as single channel in top panel) (magenta), phalloidin (grey) and DAPI (blue) with mock or SARS-CoV-2 infection at 72h after infection. Scalebar 50 μm. **F)** Quantification of dsRNA staining intensity in Calu-3 and HMVEC-L. n=9 images from 3 independent experiments. **G)** qPCR shows presence of viral RNA (MPRO) in SARS-CoV-2 infected Calu-3 and HMVEC-L co-cultured cells represented as fold change relative to mock infection at 72h after infection. n=3 independent experiments. **H)** Viral replication shown as number of PFU per ml of supernatant from SARS-CoV-2 infected Calu-3 and HMVEC-L co-cultured cells at 72h after infection. n=3 independent experiments. Measurement of cytokines with AlphaLISA Immunoassay kit for **I)** CXCL10 and **J)** IL6 in the supernatant of Calu-3 and HMVEC-L co-cultured cells with mock or SARS-CoV-2 infection at 72h after infection. n=4 independent experiments. Data are presented as mean±s.e.m. with individual data points indicated and colour coded per independent experimental replicate. Statistical significance was determined using Mann-Whitney test between mock and + SARS-CoV-2 (B, D, F, G, H, I, J). *P<0.05, **P<0.01, ****P<0.0001.

## DISCUSSION

SARS-CoV-2 can induce vascular dysfunction and thrombotic events in patients with severe COVID-19. However, the cellular and molecular mechanisms by which SARS-CoV-2 infection triggers such pathologies have so far remained elusive. Here, we have shown that endothelial cells do not support productive replication of SARS-CoV-2, but that they induce pro-inflammatory cytokines and adhesion molecules upon either direct or indirect exposure to SARS-CoV-2 virions. This supports the hypothesis that the endothelial dysfunction observed in COVID-19 patients is likely to be largely mediated by inflammatory signalling pathways.

As COVID-19 case numbers rose, reports of COVID-19 coagulopathy and vascular dysfunction also began to accumulate. Whether SARS-CoV-2 directly infects the vasculature to drive this vascular pathology has thus remained a topic of intense debate. Multiple studies show viral particles surrounding the vasculature [3, 25, 43], yet it is unclear whether this infection is truly specific to the endothelium or occurs within the perivascular compartment. Our *in vivo* analyses of lung tissue samples from deceased COVID-19 patients indicates the endothelium remains uninfected. Nevertheless, we showed that *in vitro*, endothelial cells are theoretically susceptible to SARS-CoV-2 infection, as they express ACE2, TMPRSS2 and NRP1, albeit at significantly lower levels than epithelial cells. This is in line with other studies that showed expression of SARS-CoV-2 receptors by the endothelium in human tissue or in cultured cells ^14, 17, 23, 31^. However, we did not detect endothelial cell infection following exposure to 6 x 10^4^ PFUs of SARS-CoV-2, whereas epithelial cells were readily infected at the same viral dose. These results support those of Wang et al. ^31^, who showed that epithelial cells are more susceptible to infection than endothelial cells. When exposed to higher viral titres, endothelial cells were positive for viral NP protein, suggesting effective viral entry. However, no infectious virions were detected in the endothelial cell supernatant, indicating that this infection was abortive. It is interesting to speculate as to why SARS-CoV-2 infection is productive in a monoculture of epithelial, but not endothelial cells. This may relate to cell-dependent differences in viral entry. For example, in epithelial cells, there may be sufficient surface TMPRSS2 levels available to cleave the spike protein at the S2’ site and mediate viral-host cell membrane fusion. This may in turn allow for efficient release of viral RNA into the cytoplasm and thus viral replication. In contrast, in cells with lower TMPRSS2 expression (such as endothelial cells), SARS-CoV-2 may enter the cell via endocytosis, with S2’ cleavage being mediated by endosomal cathepsins ^12^. Endosomal entry may not only be inefficient, but it may also activate the cellular anti-viral response and thereby limit productive viral replication ^48, 49^. Alternatively, a host of other cellular co-factors expressed preferentially in epithelial cells may support selective SARS-CoV-2 replication, or endothelial-specific restriction factors may limit effective viral replication.

In contrast to endothelial cells in a monoculture, viral proteins could not be detected in endothelial cells grown in a co-culture with SARS-CoV-2-positive epithelial cells, despite being exposed to the same dose of virus. These data may reflect the fact that SARS-CoV-2 typically infects and buds from epithelial cells in an apical manner, resulting in limited exposure of the endothelium to infectious virions. Extrapolating these data to the *in vivo* situation would suggest that a basolateral (abortive) infection of the endothelium by SARS-CoV-2 is only likely to occur when the epithelial barrier of the lung is severely damaged, exposing the endothelium to incoming virions.

While endothelial cells (either in a monoculture or in a co-culture) did not produce infectious virions, they did respond to SARS-CoV-2 infection. Analysis of infected endothelial monolayers revealed that cells positive for viral proteins were extruded in an apical fashion, without any apparent disruption to the monolayer, as assessed by phalloidin expression. This is in contrast to work from Buzhdygan and colleagues, who demonstrated that purified SARS-CoV-2 spike protein can induce the breakdown of the vascular barrier in brain microvascular endothelial cells ^50^. Discrepancies between these results may be due to different sources of primary endothelial cells (lung versus brain), or the differences in response of the endothelium to a purified viral protein vs infectious virus ^1, 3, 24, 29, 51–53^. However, if infected cells are promptly removed from exposed vessels as our data suggests, then this may explain why detection of viral particles in the endothelium of patients has varied between studies ^4, 54^.

In the absence of direct endothelial infection, hyperinflammation is likely to contribute to endothelial dysfunction observed in COVID-19 patients. SARS-CoV-2 triggered an upregulation of endothelial ICAM-1 expression when HMVEC-Ls were directly exposed to virus, or co-cultured with infected epithelial cells. ICAM-1 enables immune cells to effectively extravasate into tissues, so its upregulation is in keeping with the observed influx of inflammatory monocytes, neutrophils and other immune cells into the lungs in severe COVID-19 patients ^55^. This suggests that locally produced pro-inflammatory cytokines stimulate the endothelium to induce adhesion marker expression. Our previous studies with influenza virus showed that endothelial cells can themselves be a source of these pro-inflammatory cytokines ^56^, and indeed we also observed SARS-CoV-2-induced release of IL-6 and CXCL10 in cocultures. Precisely how SARS-CoV-2 is sensed by the endothelium in either the monoculture or coculture system remains to be determined. Given that SARS-CoV-2 does not replicate in endothelial cells, we speculate that endothelial cells are unlikely to sense this virus through cytosolic RNA receptors such as RIG-I or MDA5, as these immune sensors would detect viral replication intermediates. A more likely scenario is that the endothelial cells respond to danger signals from neighbouring infected epithelial or endothelial cells. These data add to a growing body of literature that indicate that endothelial cell-directed inflammation plays a key role in the pathogenesis of SARS-CoV-2 ^57^. It is tempting to speculate that, similar to influenza virus, the SARS-CoV-2-induced pro-inflammatory state may induce endothelial cells to express tissue factor, which in turn induces a pro-coagulant state, microvascular leakage and pulmonary haemorrhage ^58^, all of which have been described in patients with severe COVID-19. Furthermore, regardless of direct or indirect infection, elevated IL-6 correlates with increased fibrinogen ^59^ and although still controversial, elevated fibrinogen levels have been detected in critically ill patients ^2^.

The present study was subject to several limitations. Firstly, coagulation, thrombosis and induction of angiogenesis in the lungs of deceased COVID-19 patients has been described in addition to endothelial dysfunction ^3^. This angiogenic response is thought to be predominantly mediated through intussusceptive angiogenesis, where a new blood vessel is formed by splitting of an existing vessel. This effect appears to be specific to COVID-19 patients, as lungs from decreased influenza patients do not display increased angiogenic features. Whether this is due to relative hypoxia in the lungs remains unclear, although elevated angiogenic growth factors such as VEGF-A and VEGF-C have been associated with COVID-19 ^3, 44^ In this study we have not addressed the effect of SARS-CoV-2 exposure on the angiogenic capacity of endothelial cells in culture. Given the intimate association between inflammation, endothelial dysfunction and angiogenesis ^45, 60^, these are critical aspects of COVID-19 pathology that remain to be addressed in subsequent studies. Similarly, due to the very low levels of ACE2 present on endothelial cells it was not possible to determine if this was more pronounced on the apical or basolateral surface of the endothelial cell.

An additional limitation of the present study is that, due to the reductionist nature of the *in vitro* systems used herein, we were unable to address the contribution of immune cells to endothelial cell dysfunction during SARS-CoV-2 infection. Multiple studies attribute the sustained inflammatory response directly to immune cells, wherein macrophage activation, monocyte NLRP3 inflammasome signalling, complement activation and extrusion of neutrophil extracellular traps are implicated ^46, 61–63^. Accordingly, our data may best reflect the initial stages of infection when only a limited number of tissue-resident leukocytes may be present. Future studies will necessitate the addition of leukocytes to our co-culture system, to determine whether their presence results in a further induction of inflammation and markers of severe disease.

Here, we have conclusively shown that *in vitro*, endothelial cells are not productively infected by SARS-CoV-2 but that they mount an inflammatory response after direct or indirect exposure to the virus, characterised by increased cytokine secretion and expression of adhesion molecules. Our results thus suggest a key role of the endothelium in the pathogenesis of COVID-19, and that targeting the inflammatory response may present the best opportunity to prevent endothelial dysfunction.

## ACKNOWLEDGEMENTS AND FUNDING STATEMENT

We thank Dr. Fernando Guimaraes for assisting with acquiring samples necessary for this study. This work is supported by the National Health and Medical Research Council of Australia (Fellowship 1141131 to KS, Fellowship 1124162 to LL, Project Grant APP1158002 to EG) and the Australian Research Council (Fellowship DE180100512 to KRS, Fellowship DE170100167 to EG, Discovery Project DP200100737 to EG and AL), The National Heart Foundation of Australia (Future Leader Fellowship 104692 to EG) and UQ Early Career Researcher Grants (UQECR2058733 to LS, UQECR2058045 to LL). Imaging was performed in IMB Microscopy incorporating the Cancer Ultrastructure and Function Facility funded by the Australian Cancer Research Foundation.

## DATA AVAILABILITY STATEMENT

All raw data is available upon request.

---

**Supplemental Table 1.** Patient information. Additional information on cohort of COVID-19 patients. *representative images derived from this patient

| Patient ID | Diagnosis | Sex | Age | Co-morbidities | Histology | D-Dimer at admission |
| --- | --- | --- | --- | --- | --- | --- |
| LN1* | COVID-19 | F | 87 | Systemic Arterial Hypertension, Dyslipidemia, Hypothyroidism, Senile dementia | Acute phase DAD | 2014 |
| LN2 | COVID-19 | M | 53 | Class II obesity | Acute phase DAD | 425 |
| LN3 | COVID-19 | F | 85 | Systemic Arterial Hypertension, Brain Stroke, Vascular Dementia | Acute phase DAD | N/A |
| LN4 | COVID-19 | M | 73 | Type 2 Diabetes Mellitus, Chronic Kidney Disease Dialysis, Atrial Fibrillation, Coronary Disease, Heart Failure, Peripheral Obstructive Artery Disease | Reactive Type 2 pneumocyte hyperplasia | 3436 |
| LN6 | COVID-19 | M | 80 | Systemic Arterial Hypertension, Coronary Disease, Heart Failure, Class III obesity | Reactive Type 2 pneumocyte hyperplasia | 816 |
| LN7 | COVID-19 | M | 81 | Systemic Arterial Hypertension, Chronic Kidney Disease Dialysis, Brain Stroke | Reactive Type 2 pneumocyte hyperplasia | 13662 |
| LN8 | COVID-19 | N/A | N/A | N/A | Organising pneumonia | N/A |
| LN9 | COVID-19 | M | 86 | Prostate cancer, Abdominal Aortic Aneurysm, Giant Cells Arteritis | Organising pneumonia | 11184 |
| LN10 | COVID-19 | M | 46 | Dyslipidemia | Organising pneumonia | 594 |
| LN11 | COVID-19 | F | 93 | Type 2 Diabetes Mellitus, Systemic Arterial Hypertension, Dyslipidemia, Senile dementia | Acute phase DAD | 6544 |

**Supplemental Figure 1.**
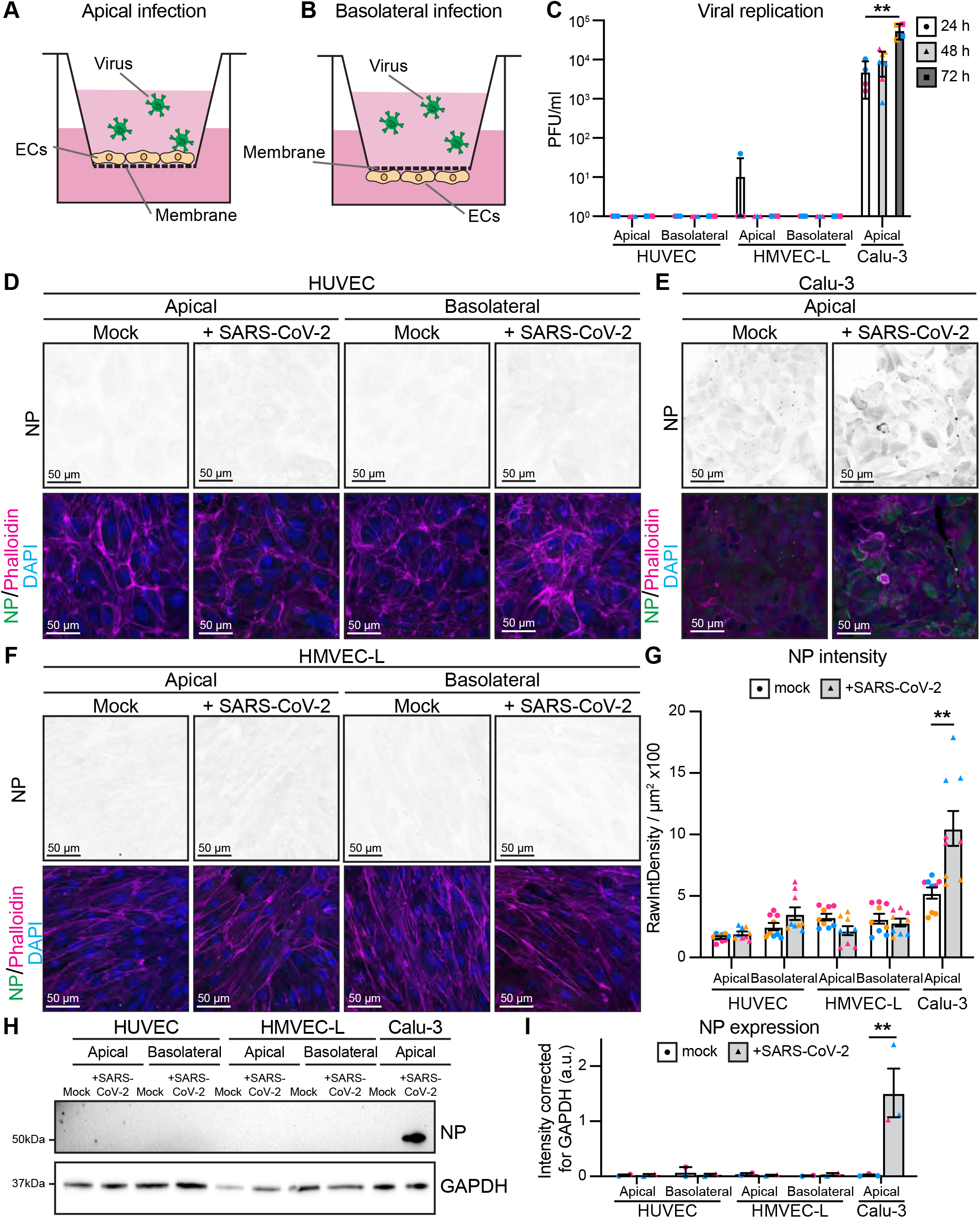
Endothelial cells are not productively infected with 6 x 10^4^ PFU of SARS-CoV-2. Schematic of **A)** apical and **B)** basolateral infection of cells cultured on transwell membranes. **C)** Viral replication shown as number of PFU per ml of supernatant from SARS-CoV-2 infected HUVEC, HMVEC-L and Calu-3 cells at 24h, 48h and 72h after infection. n=2 (HUVEC and HMVEC-L), n=3 (Calu-3) independent experiments. Representative immunofluorescent images of **D)** HUVEC, **E)** Calu-3 or **F)** HMVEC-L stained for NP (shown as single channel in top panel) (green), phalloidin (magenta) and DAPI (blue) with mock or SARS-CoV-2 infection from either apical or basolateral side of the cells at 48h after infection. Scalebar 50 μm. **G)** Quantification of NP staining intensity in HUVEC, HMVEC-L and Calu-3. n=9 images from 3 independent experiments. **H)** Western blot analysis showing NP protein levels in HUVEC, HMVEC-L and Calu-3 cells after 48h of infection. **I)** Quantification of protein levels for NP in HUVEC, HMVEC-L and Calu-3. n=2 independent experiments. Data are presented as mean±s.e.m. with individual data points indicated and colour coded per independent experimental replicate. Statistical significance was determined using Kruskal Wallis test between 24h and other time points (C) or Mann-Whitney test between mock and + SARS-CoV-2 (G, I). *P<0.05, **P<0.01

**Supplemental Figure 2.**
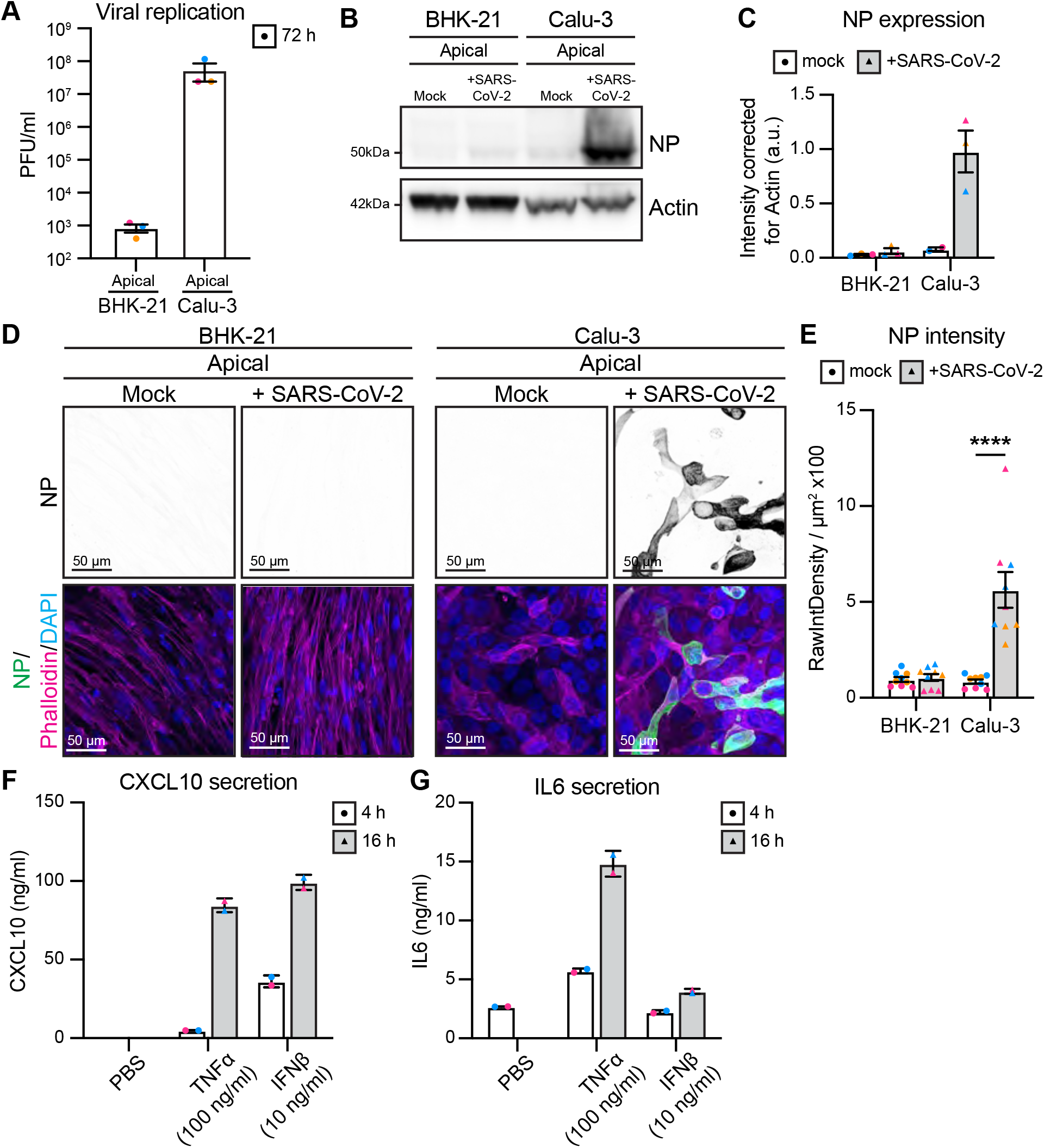
Positive and negative controls for infection with 6 x 10^4^ PFU of SARS-CoV-2. **A)** Viral replication shown as number of PFU per ml of supernatant from SARS-CoV-2 infected BHK-2 and Calu-3 cells at 72h after infection. n=3 independent experiments. **B)** Western blot analysis showing NP protein levels in BHK-21 and Calu-3 cells after 72h of apical infection with mock or SARS-CoV-2. **C)** Quantification of protein levels for NP in BHK-21 and Calu-3. n=3 independent experiment. **D)** Representative immunofluorescent images of BHK-21 and Calu-3 cells stained for NP (shown as single channel in top panel) (green), phalloidin (magenta) and DAPI (blue) with mock or SARS-CoV-2 infection from apical side of the cells at 72h after infection. Scalebar 50 μm. **E)** Quantification of NP staining intensity in BHK-21 and Calu-3. n=9 images from 3 independent experiments. Maximum secretion levels of **F)** CXCL10 and **G)** IL6 in the supernatant of HMVEC-L with mock, TNFα or IFNß treatment from the apical side of the cells at 4h and 16h after treatment. n=2 independent experiments. Data are presented as mean±s.e.m. with individual data points indicated and colour coded per independent experimental replicate. Statistical significance was determined using Mann-Whitney test between mock and + SARS-CoV-2 (C, E). ****P<0.0001

**Supplemental Figure 3.**
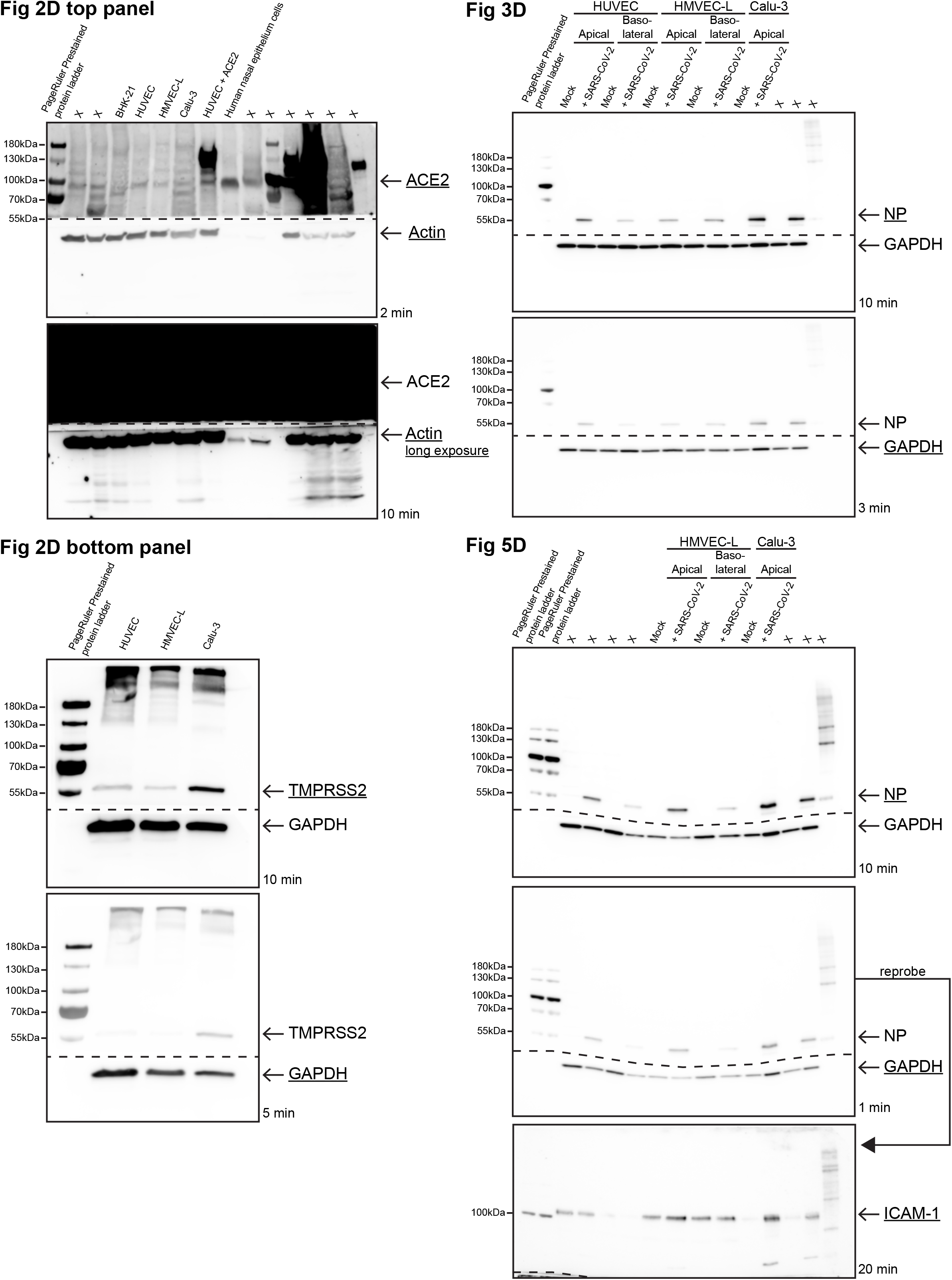
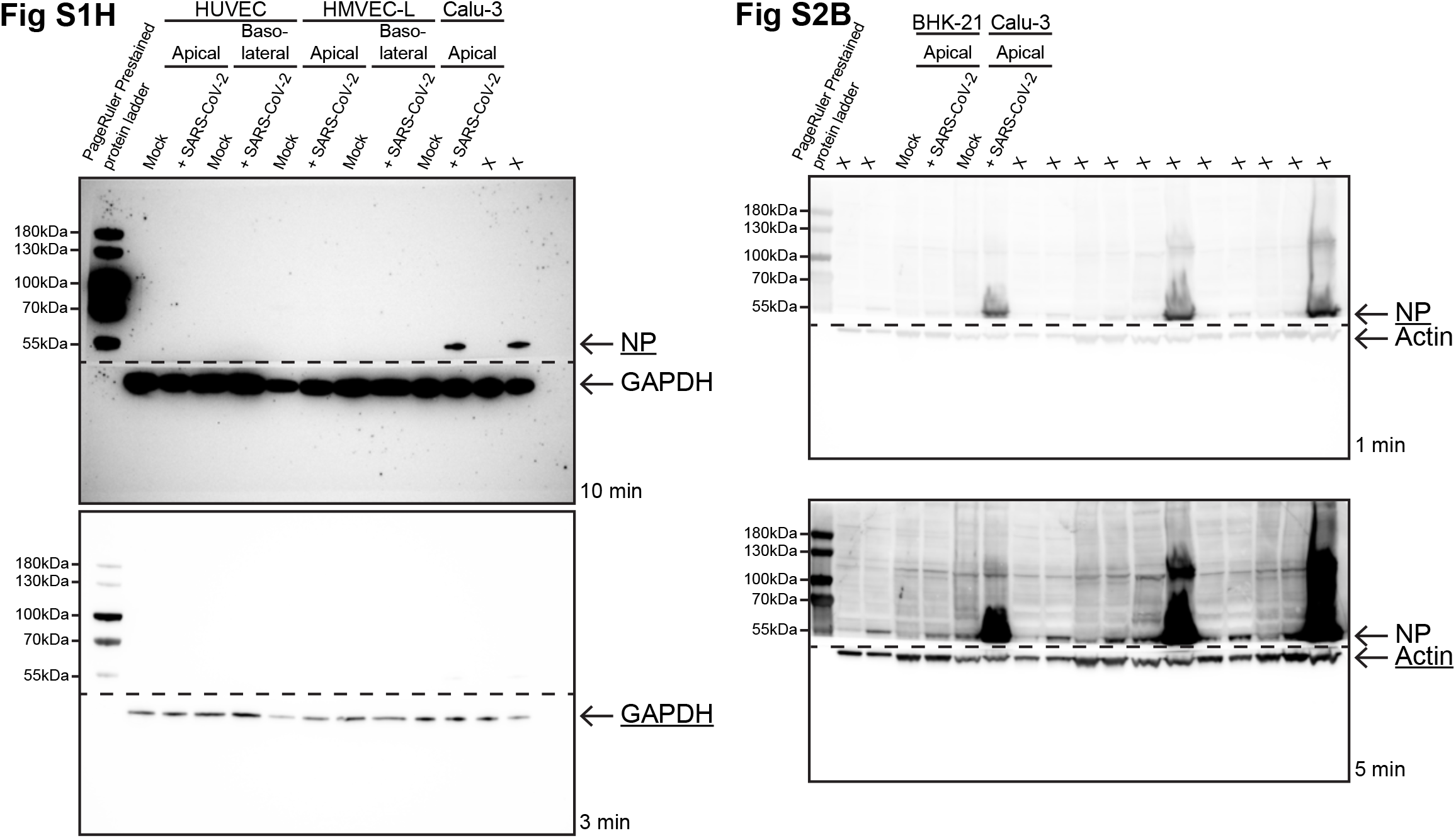
Full scans of Western blots. Full scans of Western blots shown in the paper figures. Used lanes are labelled as in the corresponding figure, with other lanes labelled as X. Dashed line indicates where membrane was cut. Underlined proteins indicate which blot was used in the corresponding figure and time indicates exposure time.

